# Understanding LRRK2 kinase activity in preclinical models and human subjects through quantitative analysis of LRRK2 and pRab10

**DOI:** 10.1101/2021.02.23.432545

**Authors:** Xiang Wang, Elvira Negrou, Michael T. Maloney, Vitaliy V Bondar, Shan V. Andrews, Manuel Montalban, Ceyda Llapashtica, Romeo Maciuca, Hoang Nguyen, Hilda Solanoy, Annie Arguello, Laralynne Przybyla, Nathan J. Moerke, Sarah Huntwork-Rodriguez, Anastasia G. Henry

## Abstract

Variants in the *leucine-rich repeat kinase 2* (*LRRK2*) gene are associated with increased risk for familial and sporadic Parkinson’s disease (PD). Pathogenic variants in LRRK2, including the common variant G2019S, result in increased LRRK2 kinase activity, supporting the therapeutic potential of LRRK2 kinase inhibitors for PD. To better understand the role of LRRK2 in disease and to support the clinical development of LRRK2 inhibitors, quantitative and high-throughput assays to measure LRRK2 levels and activity are needed. We developed and applied such assays to measure the levels of LRRK2 as well as the phosphorylation of LRRK2 itself or one of its substrates, Rab10 (pT73 Rab10). We observed increased LRRK2 activity in various cellular models of disease, including iPSC-derived microglia, as well as in human subjects carrying disease-linked variant in LRRK2 (G2019S). Capitalizing on the high-throughput and sensitive nature of these assays, we detected a significant reduction in LRRK2 activity in subjects carrying missense variants in LRRK2 associated with reduced disease risk. Finally, we optimized these assays to enable analysis of LRRK2 activity following inhibition in human peripheral blood mononuclear cells (PBMCs) and whole blood, demonstrating their potential utility as biomarkers to assess changes in LRRK2 expression and activity in the clinic.

## Introduction

Variants in the *LRRK2* gene are one of the most common genetic risk factors for Parkinson’s disease (PD) and have been shown to modify risk for both familial and sporadic forms of PD as well as immune-related diseases including Crohn’s disease (*1–3)*. LRRK2 is a multidomain protein that contains a GTPase domain, kinase domain, and several potential protein-protein interaction domains (4, *5)*. Disease-associated variants, including the most common pathogenic variant, LRRK2 G2019S, reside within the central catalytic core of LRRK2 and are reported to ultimately lead to increased LRRK2 kinase activity (6–*8)*. Further, elevated LRRK2 activity has been reported in PD patients carrying variants in other disease-linked genes or with no known genetic cause, suggesting LRRK2 may broadly contribute to the pathogenesis of PD (9, *10*).

Our understanding of how LRRK2 activity is regulated has significantly expanded in recent years through the identification of its direct physiological substrates as well as phosphorylation sites on LRRK2 that are dependent on its kinase activity. LRRK2 phosphorylates a conserved residue on the switch-II domain of a subset of Rab GTPases, including Rab10 (*11*). PD-associated variants in LRRK2 increase phosphorylation of these Rab GTPases, and elevated Rab phosphorylation likely impairs their ability to interact with downstream effectors and disrupts various aspects of intracellular trafficking, particularly in the endo-lysosomal system (*12, 13*).

LRRK2 is also constitutively phosphorylated at a set of serine residues, including Ser935, within the ankyrin and LRR regions of the protein that promotes 14-3-3 binding, and these same sites are dephosphorylated in response to LRRK2 kinase inhibition and have been utilized to assess the extent of LRRK2 inhibition in cellular and *in vivo* models (*14–16*). LRRK2 can undergo autophosphorylation at Ser1292, but analysis of phosphorylation at this site has been of limited utility given its low stoichiometry under endogenous LRRK2 expression (*17*). While analysis of phosphorylation of LRRK2 and its substrates has elucidated many aspects of how LRRK2 levels and activity are modulated, many questions remain unanswered and have been hindered by a lack of highly sensitive and quantitative assays to measure its kinase activity.

We developed high-throughput Meso Scale Discovery (MSD)-based assays to measure the phosphorylation of LRRK2 and one of its direct substrates, Rab10, and used these assays to identify novel aspects of LRRK2 regulation and to support the clinical development of LRRK2 inhibitors by enabling analysis of LRRK2 activity in accessible human samples. We demonstrated that LRRK2 is highly expressed in glial cells, including human iPSC-derived microglia, and that LRRK2 activity is significantly increased in response to lysosomal stress and inflammatory stimuli in these cells. Taking advantage of the sensitive and high-throughput nature of these assays, we were able to detect elevated LRRK2 activity in PBMCs from human subjects that carry the LRRK2 G2019S variant. Importantly, we demonstrated that the LRRK2 N551K R1398H variant associated with reduced risk for PD and Crohn’s disease leads to a reduction in Rab10 phosphorylation in both cellular models as well as in human subjects, suggesting that this variant may reduce disease risk through its effect on LRRK2’s kinase activity. Finally, we used these assays to assess LRRK2 levels and activity in whole blood and PBMCs following LRRK2 kinase inhibition, supporting their use as biomarkers to assess target and pathway engagement following dosing with LRRK2 inhibitors in the clinic.

## Results

### Development of sensitive, quantitative, and high-throughput assays to measure LRRK2 and pS935 LRRK2

We developed MSD-based assays to detect both total LRRK2 and pS935 LRRK2, a phosphorylated form of LRRK2 that is reduced following LRRK2 kinase inhibition (*14*). Antibody pairs for LRRK2 and pS935 LRRK2 were screened by ELISA assays initially using recombinant LRRK2 protein and cell lysates, and the optimal antibody pairs and orientations for capture and detection were determined based on those that gave the best dynamic range and detection specificity. Increasing levels of recombinant LRRK2 protein were assessed to determine the linearity of the assays, demonstrating a linear range of 0.16-600 ng/mL for pS935 LRRK2 and 0.06-96 ng/mL for total LRRK2 measurement and establishing the lower limits of detection (LLOD) for each assay (Fig. 1 A-B; Supplementary Table 1). To assess the specificity of the MSD-based assays in cell lysates, we used these assays to confirm LRRK2 kinase inhibition and deletion using wildtype and *LRRK2* KO A549 cells, a cell line chosen based on its high expression of LRRK2. Consistent with effects observed with western blot analysis, LRRK2 was not detected in *LRRK2* KO cells and pS935 LRRK2 levels were reduced in wildtype cells following treatment with a selective LRRK2 kinase inhibitor (MLi-2) for two hours (Fig. 1C-E). These results establish the linearity and specificity of the LRRK2 and pS935 LRRK2 MSD-based assays.

**Figure 1.**
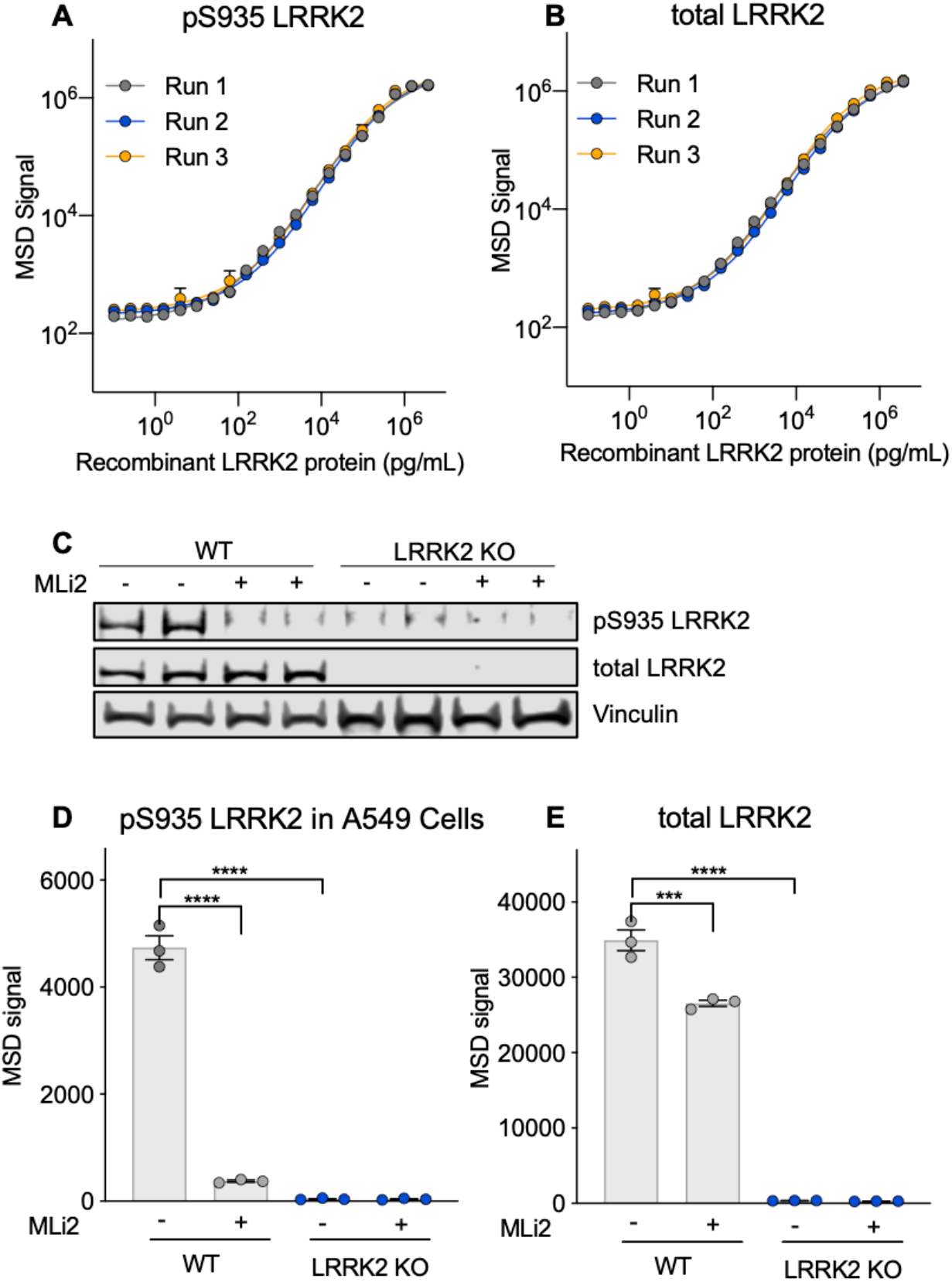
Development of specific, quantitative and high-throughput assays to measure pS935 LRRK2 and total LRRK2 **A, B**) Novel MSD assays to measure pS935 LRRK2 and total LRRK2 levels with serially diluted recombinant LRRK2 protein, n = 3. **C)** Specific detection of pS935 LRRK2 and total LRRK2 in WT and *LRRK2* KO A549 cells ± LRRK2 kinase inhibitor treatment (MLi-2, 500 nM, 2 hours) measured by western blot. **D, E**) Consistent with western blot, LRRK2 MSD assay specifically detects pS935 LRRK2 and total LRRK2 in A549 cells. n = 3. Data shown as mean ± SEM with p values: one-way ANOVA with Tukey multiple comparison test; *** p≤ 0.001, **** p ≤ 0.0001.

### Generation of a phospho-specific Rab10 antibody and development of assays to measure total and pT73 Rab10

LRRK2-dependent Rab phosphorylation has been shown to be elevated by variants in LRRK2 that increase its kinase activity and reduced following LRRK2 kinase inhibition, supporting its potential utility as a readout of LRRK2 activity (*11, 18*). Of the multiple Rabs known to undergo LRRK2-dependent phosphorylation, Rab10 has emerged as an attractive LRRK2-dependent candidate biomarker given its high expression in many cell types, including peripheral immune cells (*19, 20*). We generated a monoclonal phospho-specific antibody against pT73 Rab10, the LRRK2 phosphorylation site in Rab10, and confirmed that our antibody selectively detected phosphorylation of Rab10 and failed to detect phosphorylation of other LRRK2 Rab substrates (Fig. 2A). Using our pT73 Rab10 antibody and commercial antibodies against Rab10, we developed MSD-based assays to enable quantitative and high-throughput detection of LRRK2-dependent Rab phosphorylation as well as total levels of Rab10. The linearity and LLOD for both assays were established using recombinant human Rab10 protein that is phosphorylated at T73, with linear ranges of 65.2-8350 ng/mL and LLOD of 8.36 ng/mL for pT73 Rab10 assay and linear ranges of 0.6-45.7 ng/mL and LLOD of 0.02 ng/mL for Rab10 assay (Fig. 2B-C; Supplementary Table 2). We confirmed that these assays selectively detect LRRK2-dependent phosphorylation of Rab10 in A549 cells and matched with data obtained via western blot analysis, detecting robust reduction of pT73 Rab10 levels in wildtype cells following LRRK2 kinase inhibition and a lack of signal in *RAB10* and *LRRK2* KO cells (Fig. 2D-F; Supplementary Fig. S1). pT73 Rab10 signal was maintained in cells lacking a related LRRK2-Rab substrate (*RAB8A* KO), confirming the specificity of the assay for detecting endogenous Rab10 phosphorylation (Fig. 2D and E).

**Figure 2:**
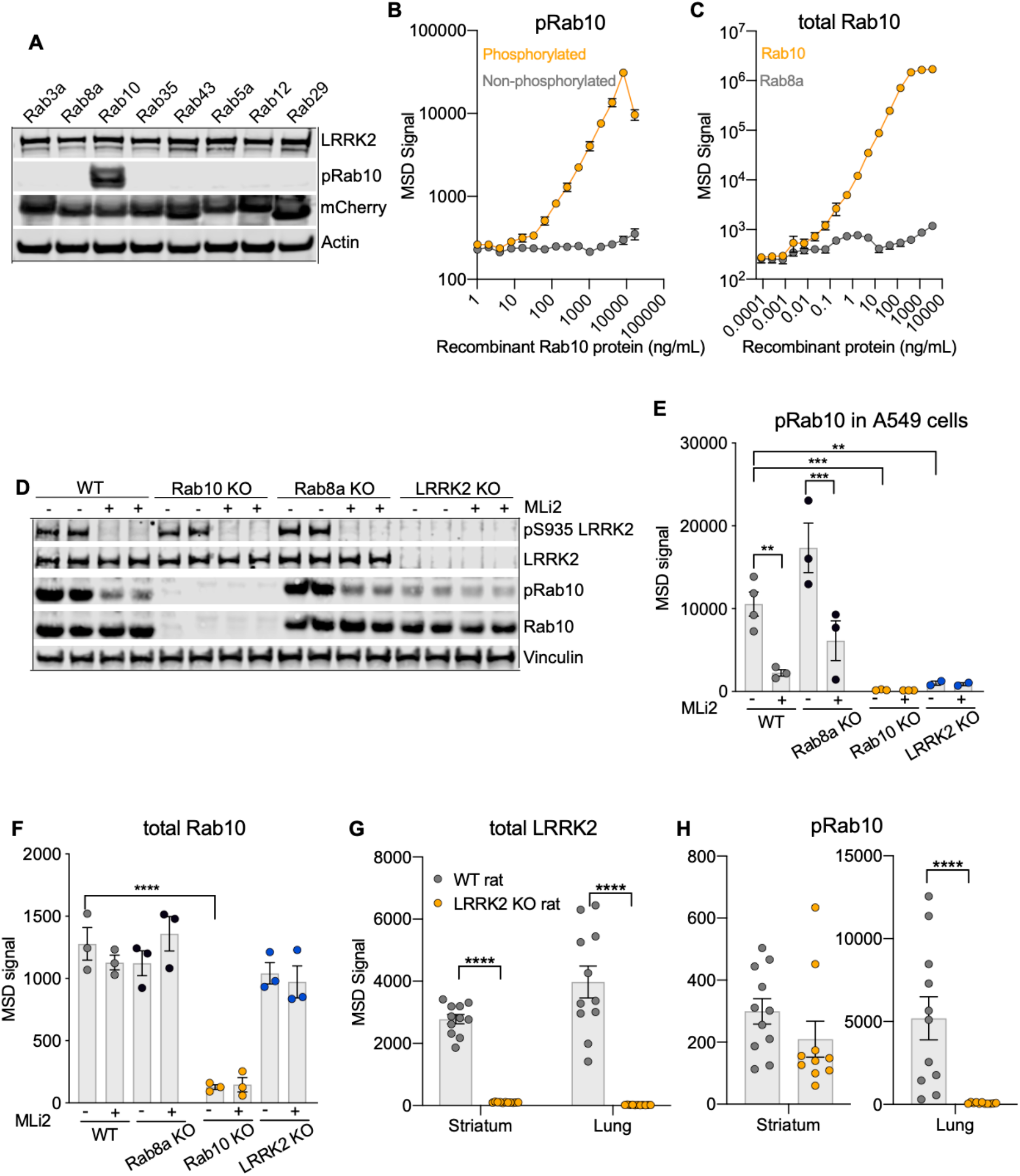
Antibody generation against pT73 Rab10 and development of MSD assays to measure pRab10 and total Rab10 in cells and *in vivo* To measure the LRRK2 kinase activity, we generated phospho-specific antibodies against the LRRK2 phosphorylation site in Rab10 (pT73 Rab10). **A**) The monoclonal pRab10 antibody DNLI 19-4 is highly selective for LRRK2 phosphorylated Rab10 in HEK293 cells overexpressing different mCherry-Rab proteins, together with FLAG-LRRK2 G2019S. **B**) A novel pRab10 MSD assay was developed to specifically measure recombinant phosphorylated Rab10 protein but not non-phosphorylated Rab10. n=3. **C)** Total Rab10 MSD assay specifically measures recombinant Rab10 protein, but not Rab8a protein. n=3. **D**) Specific detection of LRRK2 kinase dependent phosphorylation of pRab10 in wildtype, *RAB8A* KO, *RAB10* KO and *LRRK2* KO A549 cells w/wo LRRK2 kinase inhibitor MLi-2 (500 nM, 2 hours) by western blot. **E-F)** pRab10 and total Rab10 MSD assays specifically measure LRRK2-dependent phosphorylation of Rab10 and total Rab10, respectively in A549 cells. n=3. Data shown as mean ± SEM with p values: one-way ANOVA with Sidak’s multiple comparison test. **G)** Loss of pS935 LRRK2 and total LRRK2 signals in both striatum and lung from *LRRK2* KO rats. n= 11 animals for each group. **H)** *LRRK2* KO rats showed only a 30% reduction of pRab10 in striatum, and complete loss of pRab10 signals in lung. n=11 animals for each group. Direct comparison of MSD values between striatum and lung may not be valid, given that different lysis buffers were used for tissue lysates. Data shown as mean ± SEM with p values: two-way ANOVA with Sidak’s multiple comparison test; ** p≤ 0.01 *** p≤ 0.001, **** p ≤ 0.0001.

We next wanted to assess the extent of LRRK2-dependent phosphorylation of Rab10 in the brain and periphery *in vivo*. The levels of pT73 and total Rab10 were measured using our MSD-based assays in the striatum and lung of wildtype and *LRRK2* KO rats, tissues selected based on their abundant LRRK2 expression (*21, 22*). Rab10 phosphorylation was abolished in the lung of *LRRK2* KO rats compared to wildtype animals, demonstrating that phosphorylation at T73 is mediated by LRRK2 in lung (Fig. 2 G and H). pT73 Rab10 levels were reduced by approximately 30% in brain with LRRK2 deletion, suggesting that other kinase(s) may phosphorylate Rab10 in brain in addition to LRRK2 (Fig. 2H). Together, these data demonstrate that our MSD-based assays support the selective detection of LRRK2-dependent phosphorylation of Rab10 in cells and *in vivo*.

### High LRRK2 expression and activity are observed in mouse glia cells and human iPSC-derived microglia cells

While LRRK2 has been previously shown to be highly expressed in immune cells, it remains unclear which specific cell type(s) within the brain show highest levels of LRRK2 activity. To assess this, we measured the levels of LRRK2 and pT73 Rab10 in primary cortical neurons, astrocytes, and microglia cultured from wildtype mice. Consistent with previous data, LRRK2 levels were highest in mouse astrocytes and microglia and lower levels were observed in primary cortical neurons (Supplementary Fig. S2A) (*23*). Glial cells also showed the highest LRRK2 activity, with mouse astrocytes showing the highest levels of pT73 Rab10 (Fig. 3A). Treatment with a selective LRRK2 kinase inhibitor, MLi-2, significantly reduced levels of pT73 Rab10 across these CNS cell types, suggesting that phosphorylation at T73 of Rab10 in these cells is largely LRRK2 dependent. We next assessed the extent to which endogenous expression of the LRRK2 G2019S variant modulated Rab10 phosphorylation in CNS cells, focusing on cultured astrocytes from LRRK2 G2019S knock-in (KI) mice given their abundant levels of LRRK2 and pT73 Rab10. pT73 Rab10 levels were increased by nearly two-fold in primary astrocytes from LRRK2 G2019S KI mice compared to wildtype controls, demonstrating that endogenous expression of the LRRK2 G2019S variant increases Rab phosphorylation in this cell type (Fig. 3B). While total LRRK2 levels were comparable between wildtype and LRRK2 G2019S cells, we observed a significant reduction in pS935 LRRK2 levels.

**Figure 3:**
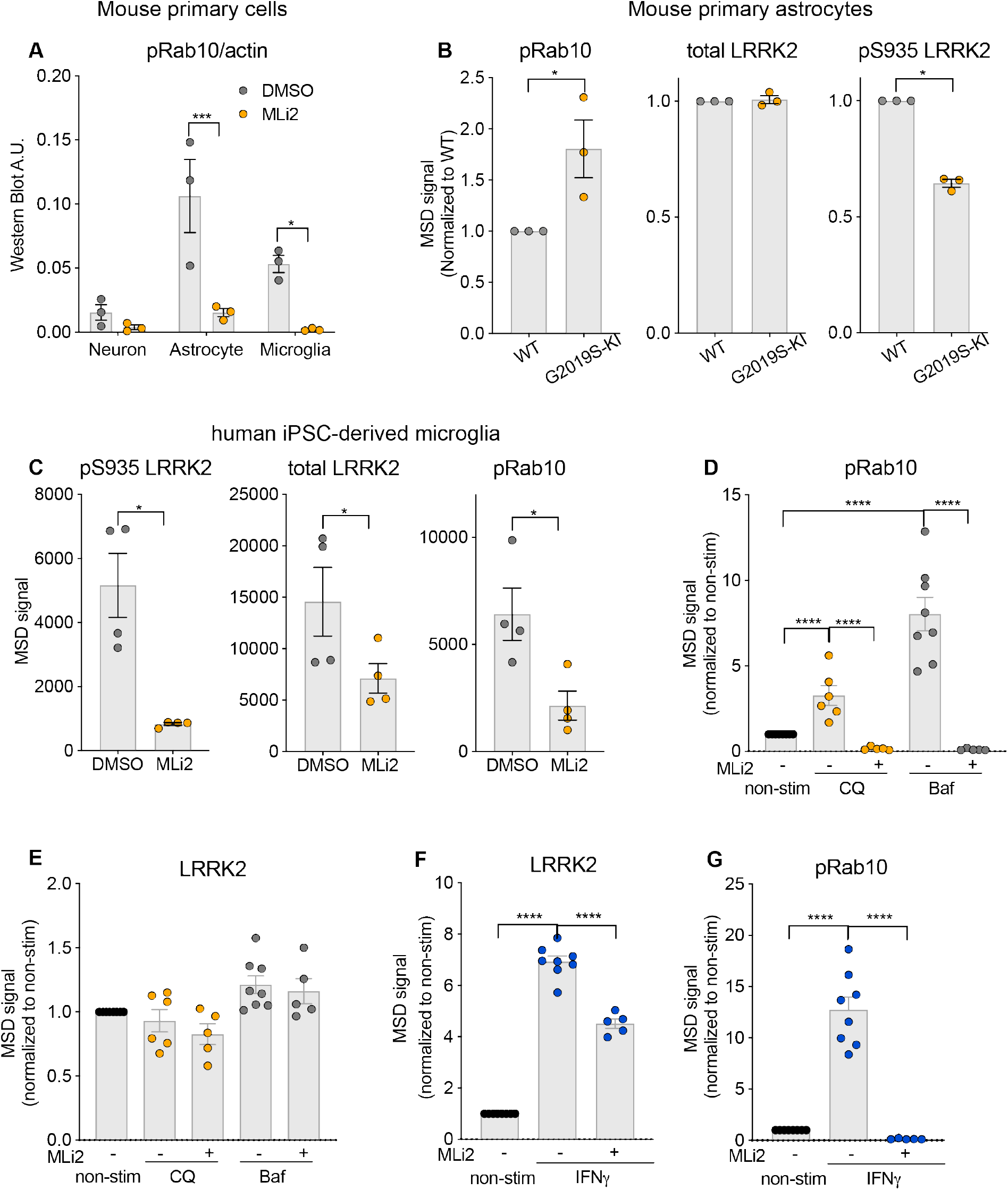
LRRK2 is highly expressed in mouse glia cells and human iPSC-derived microglia cells **A)** pRab10 is highly expressed in primary cultured mouse cortical astrocytes, and inhibition of LRRK2 kinase by MLi-2 (500 nM, 2 hours) significantly reduced phosphorylation of Rab10 in the primary cultured cortical brain cells. pRab10 levels were assessed by western blot. n=3. Data shown as mean ± SEM with p values: two-way ANOVA with Sidak’s multiple comparison test. **B)** pRab10 levels were increased in primary astrocytes from G2019S KI mice, and pS935 LRRK2 levels were decreased in mouse G2019S KI astrocytes. n=3. Data shown as mean ± SEM with p values: paired t test. **C)** Human iPSC-derived microglia expressed high levels of LRRK2 and pRab10, and LRRK2 inhibition (MLi-2, 500 nM, 2 hours) significantly reduced pRab10 levels. n=4, Data shown as mean ± SEM with p values: paired t test. **D-E)** In iMicroglia, lysosomal stresses induced by chloroquine (50 mM, 24 hours) and bafilomycin A1 (100 nM, 24 hours) resulted in increase in pRab10 levels, but not in total LRRK2 levels. **F-G)** IFN-γ treatment (20 ng/mL, 24 hours) induced a significant increase in both LRRK2 and pRab10 levels in iMicroglia cells. n=5-8. Data shown as mean ± SEM with p values: one-way ANOVA with Sidak’s multiple comparison test. * p≤ 0.05, **** p≤ 0.0001.

Transcriptome analysis of CNS cells from human cortex samples showed enrichment of *LRRK2* transcript in microglia, highlighting the potential of microglia as a relevant human CNS cell model to study mechanisms regulating LRRK2 expression and activity(*24*). We generated human iPSC-derived microglia (iMicroglia) using an established differentiation protocol and assessed the levels of LRRK2 and pT73 Rab10 basally and following LRRK2 inhibition (*25*). Strong MSD signal was observed for LRRK2, pT73 Rab10 and total Rab10 in human iMicroglia (Fig. 3C, Supplementary Fig. S2B). Rab10 phosphorylation was largely LRRK2-dependent in these cells, as pT73 Rab10 levels were significantly reduced after a two hour treatment with MLi-2. Interestingly, LRRK2 levels were also reduced at this time point, suggesting that degradation of LRRK2 might occur rapidly in human microglia upon LRRK2 inhibition. Taken together, these data show that LRRK2 is highly active in glial cells and suggest that Rab10 phosphorylation can serve as a useful readout of LRRK2 activity in these cell types.

While the expression and activity of LRRK2 are highly responsive to perturbations in cellular homeostasis in some cell types, it remains unclear whether such regulation occurs in the CNS. Previous studies have shown that pharmacological manipulations that induce lysosomal damage trigger LRRK2 translocation to lysosomes and lead to an increase in Rab phosphorylation (*26, 27*). Further, LRRK2 expression in immune cells is elevated in response to various inflammatory stimuli such as IFN- γ (*28–30*). We next assessed the extent to which lysosomotropic agents and inflammatory stimulation modulate LRRK2 levels and activity in human iMicroglia. iMicroglia were treated with bafilomycin A1(BafA1) or chloroquine (CQ), pharmacological agents that disrupt the lysosomal pH gradient and perturb lysosomal function, and the effects on LRRK2 and pT73 Rab10 were assessed. CQ and BafA1 increased pT73 Rab10 levels by three- and eightfold, respectively, as early as 6 hours post-treatment, and the effects on LRRK2 activity were sustained up to 24 hours after treatment (Fig. 3D-E; Supplementary Fig. S3A-B). Both lysosomotropic compounds increased Rab10 phosphorylation in a LRRK2-dependent fashion, as treatment with a LRRK2 kinase inhibitor significantly reduced pT73 Rab10 levels in iMicroglia (Fig. 3D). LRRK2 levels were minimally impacted following treatment with these lysosomotropic agents, with a slight, but significant, increase in LRRK2 levels observed after 6 hours of BafA1 treatment (Fig. 3E; Supplementary Fig. S3B). These data add to growing evidence that LRRK2 activity is significantly elevated in response to lysosomal stress and demonstrate that LRRK2-dependent responses to damage occur in CNS cells.

To understand whether LRRK2 levels or activity were altered in these cells in response to inflammatory stimuli, iMicroglia were stimulated with IFN- γ. Consistent with published data in peripheral immune cells, LRRK2 expression was dramatically elevated by approximately 7-fold in iMicroglia after 24 hour stimulation with IFN- γ (Fig. 3F). Elevation of LRRK2 translated to an approximately 13-fold increase in pT73 Rab10 levels, suggesting that IFN- γ treatment might affect both LRRK2 levels and activity (Fig. 3G). In support of a direct effect on LRRK2 activity, Rab10 phosphorylation was significantly, albeit more modestly, increased as early as 6 hours after IFN- γ stimulation with no significant effect on LRRK2 levels themselves (Supplementary Fig. S3C-D). The observed increase in pT73 Rab10 levels was dependent on LRRK2 kinase activity, as LRRK2 kinase inhibition prevented the elevation in Rab10 phosphorylation in response to IFN- γ treatment. These data demonstrate that LRRK2 levels and activity can both be elevated in response to inflammatory stimuli in human microglia and suggest that perturbed immune responses observed in PD patient brain might upregulate the LRRK2 signaling pathway.

### Analysis of LRRK2 and pT73 Rab10 as human blood-based biomarkers

We next applied these assays to enable the analysis of the levels of total and pS935 LRRK2 and total and pT73 Rab10 as peripheral biomarkers for use in human samples. Increasing amounts of PBMC lysates were tested to assess the linearity of the LRRK2 and Rab10 assays in this matrix. The linear range of detection for pS935 and total LRRK2 were from 12-3070 μg/mL of PBMC lysate (3070 μg/mL was the highest concentration tested, Fig. 4A; Supplementary Table 3). Measurements of total and pT73 Rab10 were slightly less sensitive, with a linear range of 191.9-3070 μg/mL of PBMC lysates for pT73 Rab10 and 3-12 ug/mL for total Rab10 (Fig. 4B; Supplementary Table 4). To confirm these assays could be employed to monitor LRRK2 inhibition, we treated PBMCs from three healthy donors with increasing concentrations of MLi-2 *ex vivo* and measured the effects of treatment on pS935 LRRK2 and pT73 Rab10 levels (Fig. 4C). LRRK2 inhibition reduced the levels of pS935 LRRK2 and pT73 Rab10 in a comparable dose-dependent manner, with IC50s of 3.8 nM for pS935 LRRK2 and 5.3 nM for pT73 Rab10.

**Figure 4.**
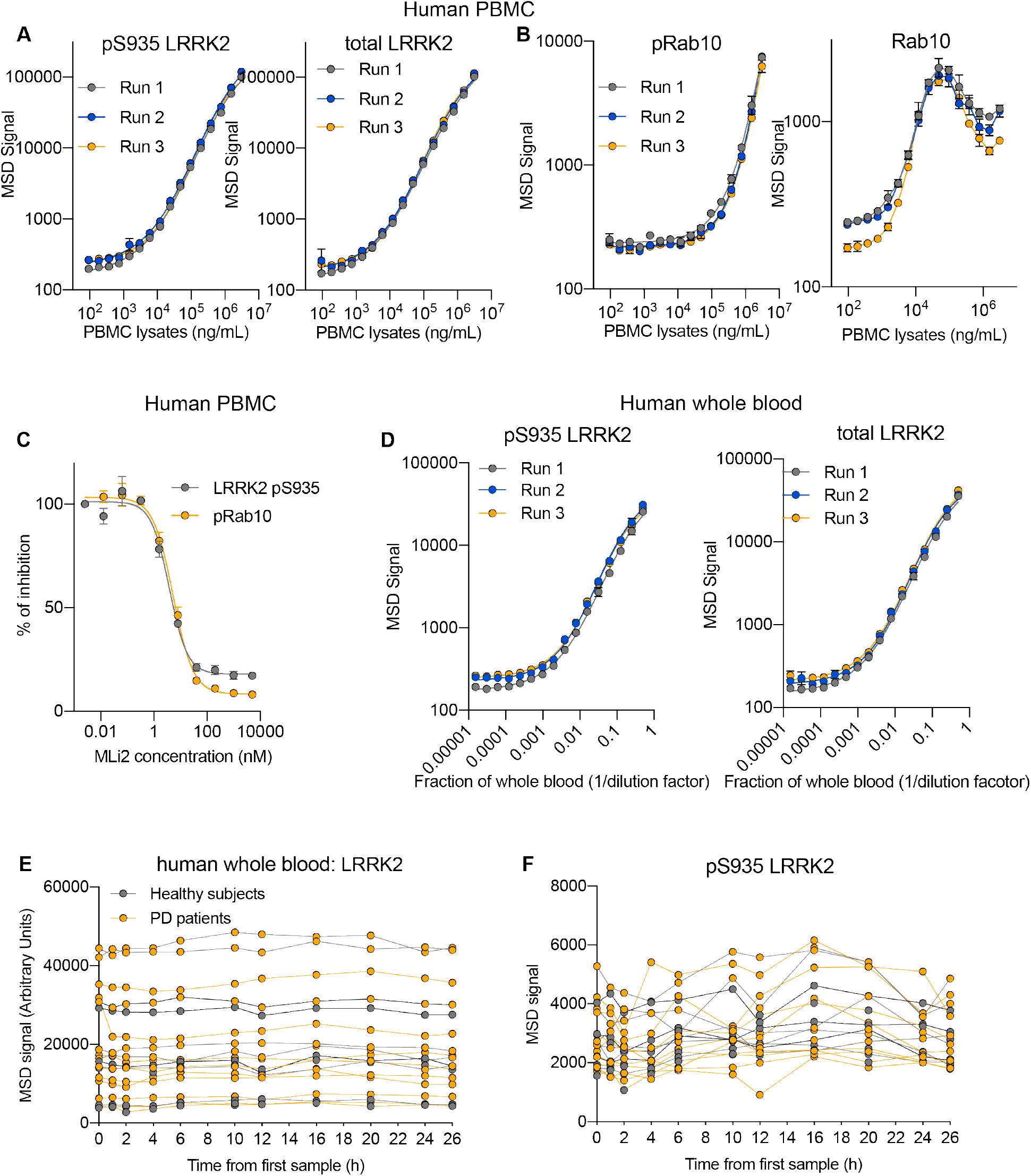
Development of LRRK2 and pRab10 assay for human blood-based biomarkers analysis. **A)** pS935 LRRK2 and total LRRK2 levels measured with serial diluted lysates from human PBMC. **B)** pRab10 and total Rab10 levels measured with serial diluted lysates from human PBMC. **C)** Inhibition of LRRK2 kinase decreased pS935 LRRK2 and pRab10 in a dose-dependent manner in PBMC with ex-vivo treatment of MLi-2 (1 hour). PBMC from n=3 donors. **D)** pS935 and total LRRK2 levels measured with serial diluted samples of human whole blood. **E)** Total LRRK2 levels in whole blood from healthy subjects and PD patients were stable within subjects over the course of 24 hours:Average coefficient of variation (CV) = 7%; average maximum to minimum value within-subject fold change of 1.25, while displaying high variability between subjects (66% CV, 11.6-fold difference in maximum to minimum value. **F)** pS935 LRRK2 levels measured in human whole blood samples varied to a greater extent than total LRRK2 within subjects: Average CV=17%, average within-subject maximum to minimum value fold change of 1.76. But between subjects the variability was less than that of total LRRK2 (36% CV, 3.8-fold difference in maximum to minimum value)

Given the sensitivity of the pS935 and LRRK2 MSD-based assays, we evaluated whether these assays could be used to directly measure LRRK2 levels and inhibition from whole blood to enable simpler analysis in a clinical setting. Blood samples from healthy donors were serially diluted, and the levels of pS935 and total LRRK2 were assessed (Fig. 4D). Both measures showed a linear increase from the range of 2 to 512 folds of dilution (Supplementary Table 5). We confirmed the specificity of detection of these assays by measuring the levels of pS935 and total LRRK2 in PBMCs and whole blood collected from wildtype or *LRRK2* KO rats following dosing with a tool LRRK2 kinase inhibitor developed by Denali, DN1388, for ten days. The pS935 LRRK2 signal was completely abolished in both PBMCs and whole blood from *LRRK2* KO rats or those dosed with 100mg/kg DN1388 (Supplementary Fig. S4). LRRK2 levels were not detected in PBMCs or whole blood from *LRRK2* KO rats. These data demonstrate the specificity of these measurements and validate the use of the whole blood assay to measure pS935 and total LRRK2.

To further assess the potential utility of LRRK2, pS935 LRRK2, and pT73 Rab10 as biomarkers in future clinical studies, we characterized the intra- and inter-subject variability of these analytes in blood. Whole blood samples from healthy controls and PD subjects was obtained from the 24-hour Biofluid Sampling Study sponsored by the Michael J Fox Foundation in which samples were collected at 11 time points over a 24-hour period, and the levels of pS935 and total LRRK2 were measured. LRRK2 levels were stable within subjects over the course of 24 hours (average coefficient of variation (CV) = 7%), while displaying high variability between subjects (66% CV) (Fig. 4E). pS935 LRRK2 levels varied to a greater extent than LRRK2 within subjects (average CV=17%) but between subjects the variability was less than that of total LRRK2 (36% CV) (Fig. 4F). The relatively low intra-subject variability and detectability of the analytes in human whole blood support the potential of these assays to assess LRRK2 levels and inhibition in blood in a clinical setting.

### LRRK2 and pT73 Rab10 levels in LRRK2 G2019S carriers compared to healthy controls

We next wanted to better define the extent to which variants in *LRRK2* associated with increased risk for developing disease affect the levels of LRRK2 and its kinase activity. Variants in LRRK2 within the ROC, COR, kinase, and WD40 domains have been linked to increase risk for PD or Crohn’s disease and have been reported to lead to increased Rab10 phosphorylation (*1, 31*). Disease-linked LRRK2 variants were expressed in HEK293T cells, and the levels of pT73 Rab10 were measured and normalized to total levels of LRRK2 to control for any differences in LRRK2 expression. Consistent with previous findings, all variants assessed led to an increase in pT73 Rab10 levels when normalized to LRRK2, with the LRRK2 G2019S variant showing an approximately threefold increase in pT73 Rab10 levels (Supplementary Fig.S5).

We exploited the high-throughput and quantitative nature of our assays to determine whether the levels or kinase activity of LRRK2 were altered in PD patients that were heterozygous for the LRRK2 G2019S variant or in those with sporadic disease. We measured LRRK2, pS935 LRRK2, and pT73 Rab10 levels in a large collection of PBMC samples obtained from healthy subjects and PD patients with and without the LRRK2 G2019S variant. pS935 LRRK2 and total LRRK2 levels were reduced in subjects with PD carrying the G2019S variant compared to sporadic PD patients and healthy LRRK2 G2019S carriers (Fig. 5A-B). This same sample set has been recently analyzed for total and pS935 LRRK2 levels via an orthogonal and analytically validated assay, with substantially similar results, as well as via a sandwich ELISA-based assay with a noted non-significant trend toward reduced pS935 LRRK2 normalized to total LRRK2 levels, providing independent validation of these results(*32, 33*). pT73 Rab10 levels were not significantly different in any group, but when normalized for variations in total LRRK2 levels, we observed a significant elevation in LRRK2 G2019S carriers with PD compared to sporadic PD patients and in healthy LRRK2 G2019S carriers compared to non-carriers (Fig. 5C-D). Further, there was a non-significant trend toward increased pRab10 normalized to total LRRK2 in sporadic PD patient (Fig. 5D). These data support that LRRK2 G2019S carriers with PD have elevated LRRK2 kinase activity that can be measured in a clinical setting.

**Figure 5.**
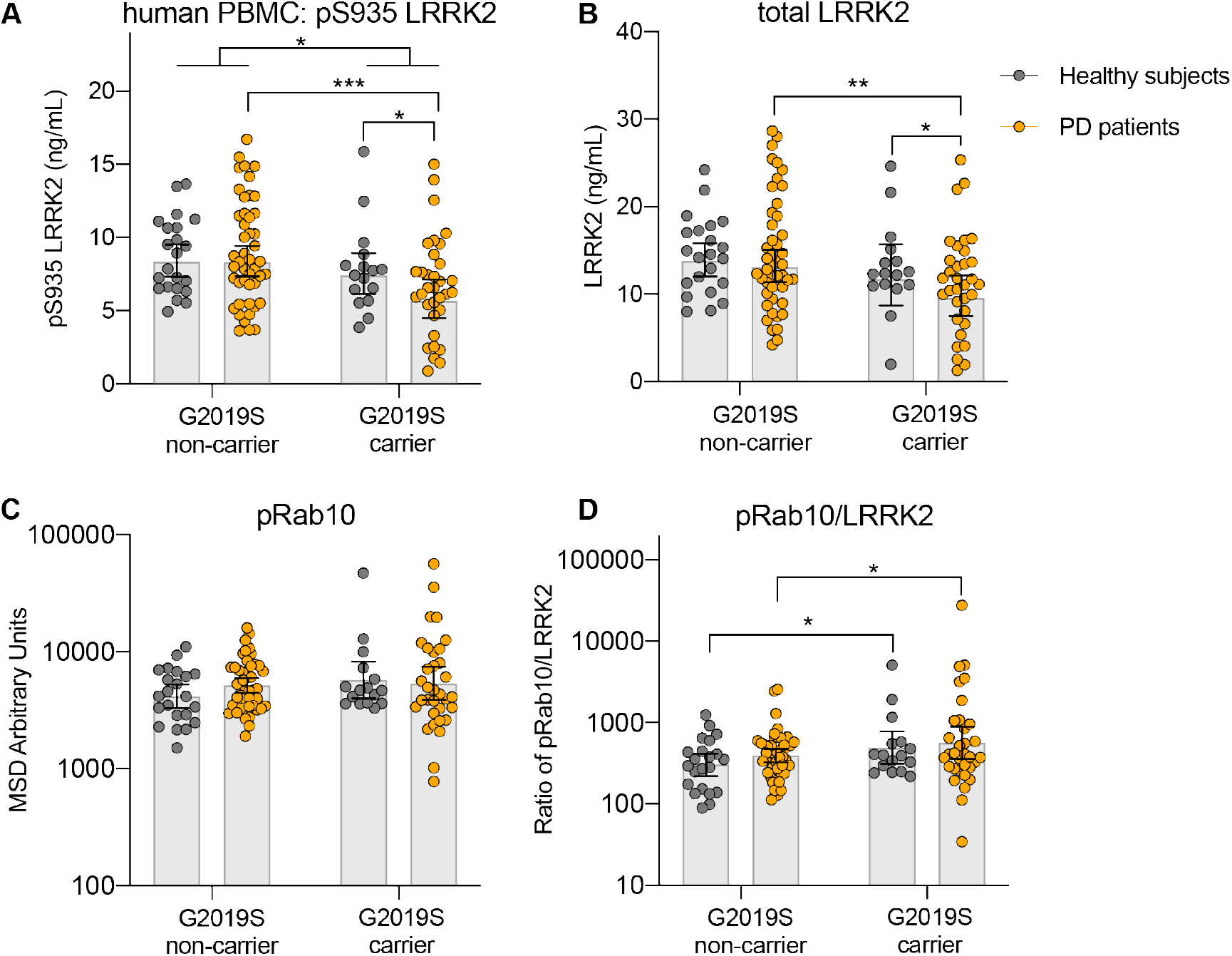
LRRK2 and pRab10 levels in PD patients and healthy subjects. LRRK2 levels and activities were measured in PBMC from PD patients and healthy subjects with and without LRRK2 G2019S variant provided by the LRRK2 Detection in PBMC Consortium (*33*). n=16 non-PD-LRRK2 G2019S carrier, n=22 non-PD non-G2019S, n=46 PD non-G2019S carrier, n=33 PD and G2019S carrier. Data shown as mean ± SEM. For each variable, an ANCOVA model was fit, with log2 (value) as response and terms for Cohort, G2019S status (and their interaction), Gender and Age. For each variable a forward model selection step was performed to assess the usefulness of adjusting for Age or including an interaction term. **A)** For pS935 LRRK2, p=0.0002 between G2019S carrier and non-carrier in PD (Relative Geometric mean ± 95% confidence interval = 0.651 (0.524 - 0.810)); p=0.0134 between PD and non-PD in G2019S carrier (Relative Geometric mean ± 95% confidence interval = 0.673 (0.492 - 0.920)); p=0.0233 between G2019S carrier and non-carrier (Relative Geometric mean ± 95% confidence interval = 0.805 (0.660 - 0.970)). **B)** For LRRK2, p=0.0028 between G2019S carrier and non-carrier in PD (Relative Geometric mean ± 95% confidence interval = 0.688 (0.540 - 0.877)); p=0.0396 between PD and non-PD in G2019S carrier (Relative Geometric mean ± 95% confidence interval = 0.693 (0.489 - 0.982)). **C-D)** For pRab10/LRRK2, p=0.0171 between G2019S carrier and non-carrier in PD and in non-PD (Relative Geometric mean ± 95% confidence interval = 1.44 (1.07 – 1.95)). Relative geometric means given are adjusted for age and sex. Data shown as geometric mean with 95% CI. *p≤ 0.05, ** p≤ 0.01, *** p≤ 0.001.

### LRRK2 N551K R1398H protective haplotype is associated with reduced pT73 Rab10 levels

We assessed whether variants in *LRRK2* associated with reduced risk for developing disease affect the levels of LRRK2 and its kinase activity. The N551K R1398H haplotype is associated with protection from developing PD and Crohn’s disease (*31, 34–36*). Previous *in vitro* studies have suggested that the R1398H variant enhances the GTPase activity of LRRK2 (*37*). However, it remains unclear how this variant affects LRRK2 activity in cells. We expressed various LRRK2 variants that possessed the N551K R1398H variant on its own or in the presence of additional variants that are associated with increased risk for PD or Crohn’s disease (Fig. 6A). Expression of the LRRK2 N551K R1398H variant on its own or together with disease associated LRRK2 variants within its kinase domain led to reduced total levels of LRRK2 (Fig. 6B). The decrease in LRRK2 levels observed with the LRRK2 N551K R1398H variant singly or together with the G2019S variant translated to a reduction in levels of pT73 Rab10 (Fig. 6C). The protective LRRK2 variants in combination with variants in the ROC, COR, or WD40 domains did not impact LRRK2 levels or activity. To confirm that the LRRK2 protective haplotype modulates LRRK2 levels and activity under endogenous expression conditions, we generated LRRK2 N551K R1398H KI A549 cells and observed a reduction in LRRK2 levels and concomitant decrease in pT73 Rab10 levels in LRRK2 N551K R1398H KI cells compared to wildtype cells (Fig. 6D-E). These data suggest that the N551K R1398H variant ultimately leads to reduced LRRK2 activity and may mediate its effects on LRRK2 stability through interactions within the GTPase or WD40 domains.

**Figure 6.**
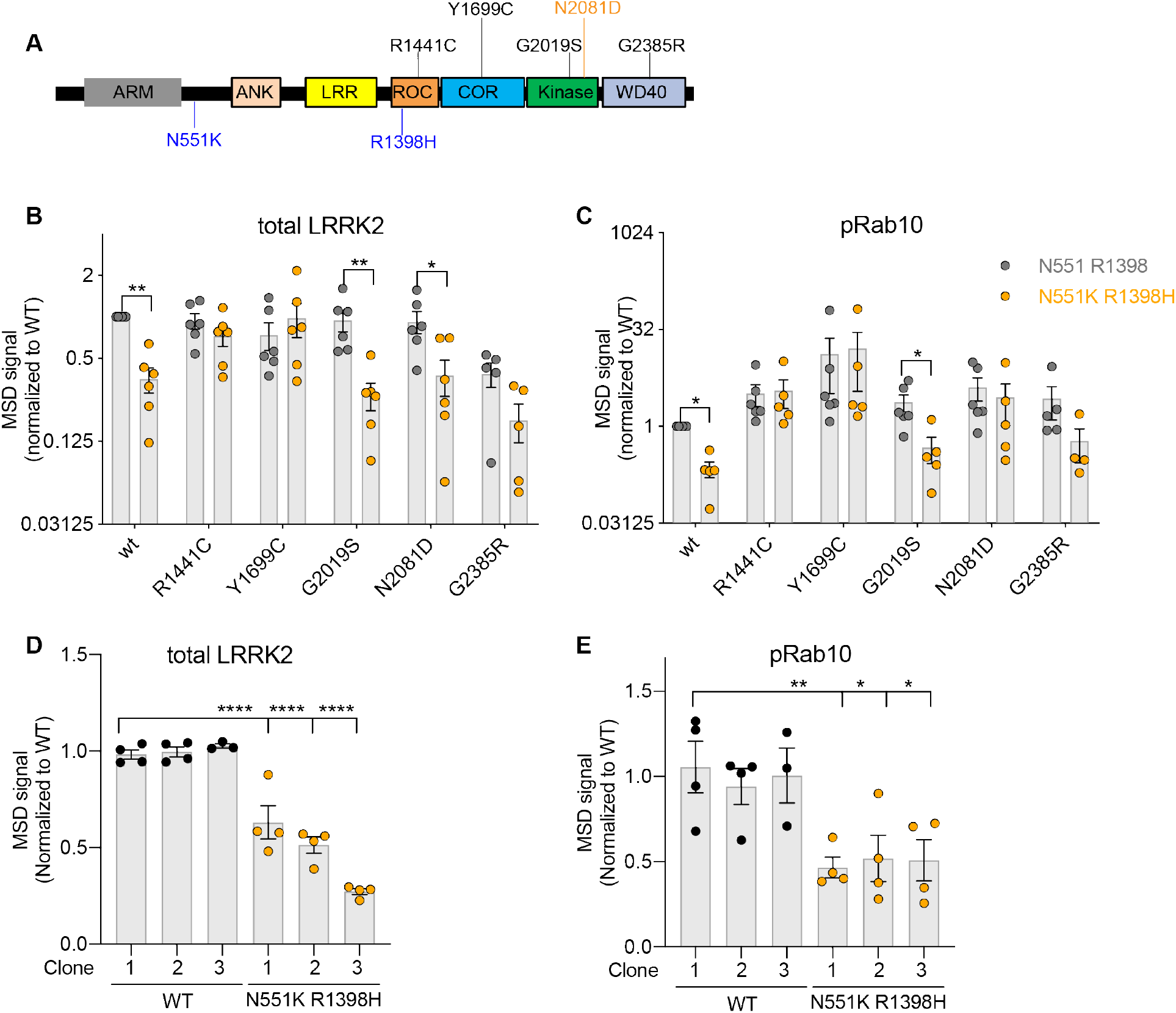
LRRK2 activity is reduced with LRRK2 protective variants associated with Parkinson’s (PD) and Crohn’s disease (CD) in cells. **A**) Schematic overview of LRRK2 risk and protective variants associated with PD and CD. N551K R1398H: protective variants associated with both PD and CD (in blue). N2081D: a risk variant associated with CD (in orange). **B-C)** In HEK293T cells overexpressing different LRRK2 variants and Rab10, total LRRK2 and pRab10 levels were reduced with LRRK2 N551K R1398H variants on its own or with PD-linked LRRK2 variants in the kinase domain. n=5-6, Data shown as mean ± SEM with p values: two-way ANOVA with Sidak’s multiple comparison test. **D-E)** LRRK2 and pRab10 levels were reduced in LRRK2 N551K R1398H KI A549 cells. Three wildtype pooled cells, and three clones of LRRK2 N551K R1398H KI A549 cells were used. n=4 batches. Data shown as mean ± SEM with p values: one-way ANOVA with Sidak’s multiple comparison test. * p≤ 0.05, ** p≤ 0.01, **** p≤ 0.0001

To expand upon our observations in cellular models and capitalize on our high-throughput assays, we assessed whether the LRRK2 protective haplotype modulates LRRK2 levels or activity in PBMCs obtained from healthy subjects and PD patients with and without the LRRK2 N551K R1398H variant. Levels of total and pS935 LRRK2 were unchanged in LRRK2 N551K R1398H carriers compared to non-carriers regardless of disease status (Fig. 7A and B). These data differ from effects observed on total LRRK2 in LRRK2 N551K R1398H cell models, suggesting the protective haplotype may affect LRRK2 levels in a cell-type dependent manner. Importantly, Rab10 phosphorylation was significantly reduced in LRRK2 N551K R1398H carriers compared to non-carriers, an effect that was maintained when normalized for total LRRK2 levels and in sensitivity analyses that additionally adjusted for disease status and/or LRRK2 G2019S status (Fig. 7C-D and Supplementary Table 6). These data provide the first evidence, to our knowledge, that the LRRK2 protective haplotype is associated with reduced LRRK2 activity in human subjects.

**Figure 7:**
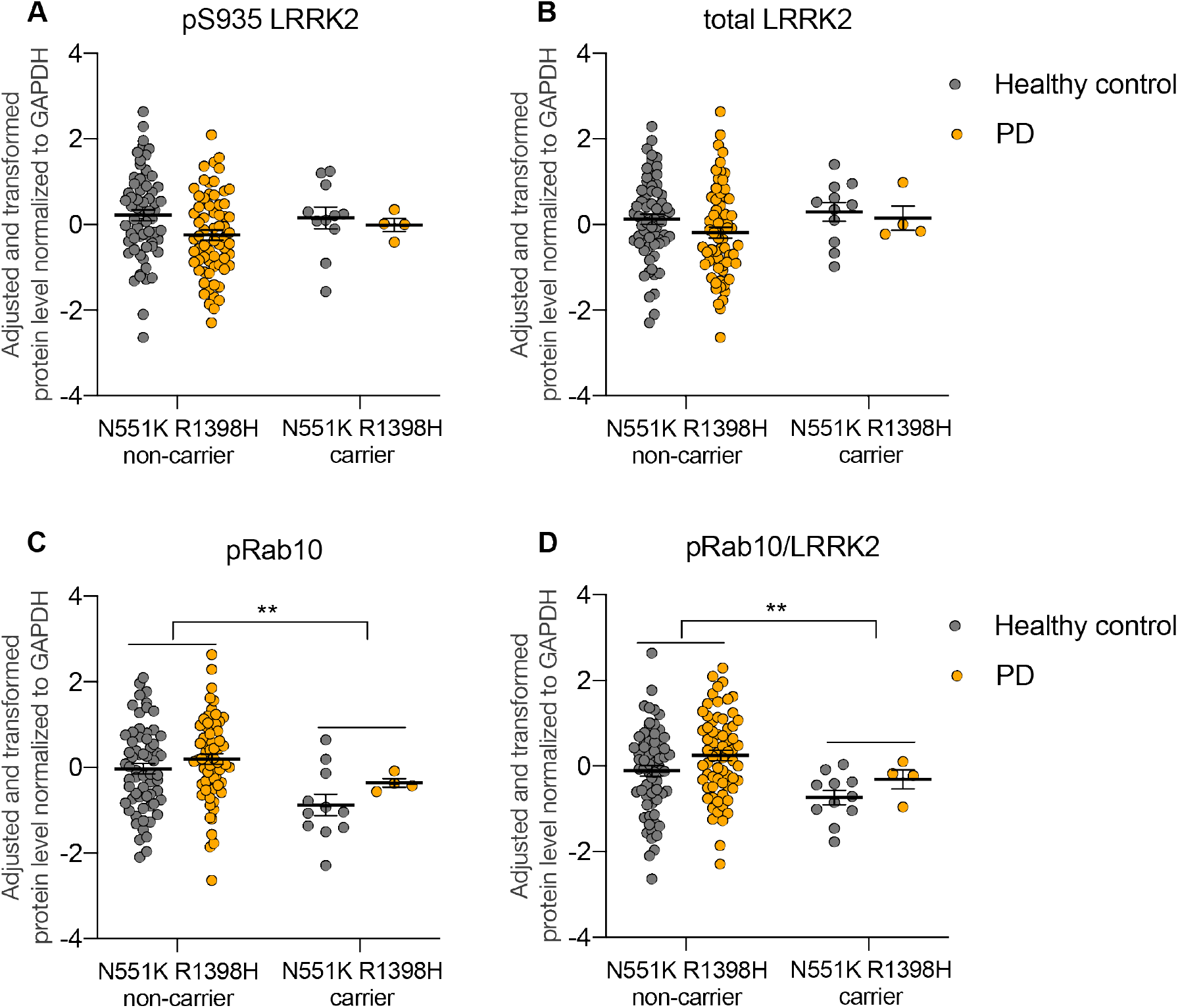
LRRK2 and pRab10 levels in N551K R1398H haplotype carriers and non-carriers. LRRK2, pS935 LRRK2 and pRab10 levels were measured in PBMCs isolated from PD and healthy subjects and N551K R1398H haplotype status was derived from genotype array data (n=64 healthy control N551K R1398H non-carriers, n=66 for PD N551K R1398H non-carriers, n=11 for healthy control N551K R1398H carriers, and n=4 for PD N551K R1398H carriers). pRab10 (0.83 standard deviation decrease in carriers vs non-carriers, p=0.0020) and the pRab10/LRRK2 ratio (0.70 standard deviation decrease in carriers vs non-carriers, p = 0.0096) were reduced in N551K R1398H carriers compared to non-carriers. pRab10 and pRab10/LRRK2 were also significantly (p < 0.05) reduced in N551K R1398H carriers relative to non-carriers in sensitivity analyses that additionally adjusted for disease status and/or LRRK2 G2019S status. Data shown as mean ± SEM. ** p≤ 0.01

## Discussion

In the current study, we describe the development of high-throughput and sensitive assays to quantify the levels and phosphorylation of LRRK2 and Rab10 and apply these assays to better understand how LRRK2 is regulated in cellular models and in human subjects that carry LRRK2 variants. Using these assays, we observed an approximately one and a half to two-fold increase in Rab10 phosphorylation with the LRRK2 G2019S variant under endogenous expression conditions in mouse astrocytes and in PBMCs from LRRK2 G2019S variant carriers after normalization for variations in levels of LRRK2. Beyond effects observed with variants in LRRK2 itself, we found Rab10 phosphorylation was significantly increased in response to pharmacological agents that perturb PD-relevant pathways in iPSC-derived microglia and demonstrated that this was dependent on the kinase activity of LRRK2. Finally, we showed that missense variants in LRRK2 associated with reduced risk for PD and Crohn’s disease lead to lower levels of Rab10 phosphorylation, suggesting that these variants may lower disease risk by reducing the activity of the LRRK2 signaling pathway. Given their successful application in human PBMCs and whole blood, these assays have the potential to enable quantitative analysis of LRRK2 levels and activity following dosing with LRRK2 inhibitors in the clinic.

An understanding of the function of LRRK2 in different cell types throughout the brain has been hindered by a lack of quantitative tools to measure LRRK2 levels and activity across CNS cell types. Previous studies have reported varying levels of LRRK2 expression in neurons and glial cells, including transcriptome analysis of neurons, glia, and vascular cells from the cortex of mouse and human samples which showed enrichment of *LRRK2* transcript in astrocytes and or microglia, respectively (*23, 24, 38–41*). We analyzed LRRK2 levels and activity across primary mouse cortical neurons, astrocytes, and microglia and confirmed high expression of LRRK2 in glial cells with weaker expression observed in neurons. Additionally, we showed that LRRK2 inhibition results in a near complete reduction in pT73 Rab10 levels across all three cell types, demonstrating that LRRK2-dependent Rab10 phosphorylation is relevant in brain. Our analysis of LRRK2 levels and activity in iPSC-derived microglia yielded similar findings and suggested that LRRK2-dependent phosphorylation of Rab10 also occurs in human microglia. Rab10 phosphorylation was not completely abolished in brain lysates from *LRRK2* KO mice, suggesting the LRRK2-dependence of Rab10 phosphorylation may be cell type or brain region specific or depend on the activation state of glial cells within the brain. Emerging data suggest that LRRK2 expression or kinase activity are modulated by inflammatory stimuli or lysosomal damage (*26, 27, 42, 43*). We demonstrated that Rab10 phosphorylation was significantly increased in iPSC-derived microglia following treatment with pharmacological agents that dissipate the endo-lysosomal pH gradient, providing support that lysosomal dysfunction can impact LRRK2 activity in human microglia and highlighting the potential of this model to better understand the mechanisms governing LRRK2 regulation. We cannot exclude the possibility that *in vitro* culture conditions may impact LRRK2 activity following lysosomal damage, and additional studies are warranted to understand whether similar regulation occurs in glial cells from PD patient brains with evidence of lysosomal dysfunction.

To enable the clinical development of LRRK2 inhibitors and analysis of alterations in LRRK2 pathway activity in PD patient populations, assays are needed that can be used to reliably quantify LRRK2 levels and phosphorylation of the kinase itself and its direct substrates in samples easily obtained from human subjects. We optimized our MSD-based assays and demonstrated linear quantification of the levels of total and pS935 LRRK2 as well as total and pT73 Rab10 in human PBMCs. *Ex vivo* treatment of PBMCs with the LRRK2 inhibitor MLi-2 resulted in dose-dependent reduction of pS935 LRRK2 and pT73 Rab10 levels with an equivalent cellular potency, confirming the utility of both readouts as measures of LRRK2 inhibition in accessible human samples. We further optimized the pS935 and total LRRK2 assays to allow the quantification of both analytes in whole blood, providing a critical advance to quantify the extent of LRRK2 inhibition reliably across multiple locations and with multiple handlers for use in a clinical setting. Future work will focus on using these assays to demonstrate target engagement and modulation of downstream pathway activity following dosing with LRRK2 inhibitors in human subjects to help assess the potential of LRRK2 inhibitors in the clinic.

While pathogenic variants in LRRK2 lead to increased kinase activity and elevated phosphorylation of its Rab substrates, the extent to which these variants, including the most common variant G2019S, increase LRRK2 kinase activity and the broader relevance of increased LRRK2 activity to sporadic PD remain controversial. Defining the degree to which LRRK2 activity is elevated in PD patients with and without variants in LRRK2 is paramount for the clinical development of LRRK2 inhibitors and can help determine how much LRRK2 inhibition should be targeted as well as better identify which patient populations might benefit most from LRRK2 inhibition. The LRRK2 G2019S variant has been reported to elevate its kinase activity by approximately two- to tenfold depending on the model system used and method employed to quantify autophosphorylation or phosphorylation of its direct substrates (*11, 17, 44–46*). As many previous studies have relied on cellular models overexpressing LRRK2 or peripheral tissues from *in vivo* models, it remains unclear the degree to which variants in LRRK2 affect its kinase activity in the CNS or the extent to which its activity is increased in PD patients. By assessing the levels of pT73 Rab10, we demonstrate that LRRK2 activity is increased nearly twofold in primary astrocytes derived from mice expressing the LRRK2 G2019S variant at endogenous levels. Further, we showed that PBMCs from heterozygous LRRK2 G2019S variant carriers also had a significant increase in Rab10 phosphorylation of approximately 1.5-fold compared to non-carriers after normalizing for variations in LRRK2 levels. Our results are largely consistent with findings from other studies that show a mild increase in LRRK2 activity in G2019S carriers compared to non-carriers, including recent data showing an approximately 1.9 fold increase in pT73 Rab10 levels in neutrophils obtained from LRRK2 G2019S carriers compared to healthy controls using a mass-spectrometry based approach(*19, 20, 47*). Together, these data add further evidence that LRRK2 G2019S activity is elevated approximately one and a half to twofold under endogenous conditions in relevant cell types, including CNS cells, and suggest a small (<2-fold) increase in phosphorylated Rab10 is sufficient to confer an elevated lifetime risk of Parkinson’s Disease. These results can serve as a useful guide to inform the level of LRRK2 kinase inhibition required to normalize LRRK2 activity in the clinic. We also observed a non-significant trend of increased Rab10 phosphorylation in PBMCs from sporadic PD patients, and additional analysis of LRRK2 activity is warranted in larger cohorts and across broader populations to better understand the range of increase in kinase activity conferred by other pathogenic LRRK2 variants or in sporadic PD more broadly.

While pathogenic variants in LRRK2 ultimately lead to an increase in its kinase activity, the functional consequences of a protective LRRK2 haplotype remained poorly understood. The N551K-R1398H-K1423K LRRK2 haplotype is associated with reduced risk for developing PD, REM behavior sleep disorder, and Crohn’s disease, with a combined odds ratio for PD of approximately 0.82 (*31, 34–36, 48*). The R1398H variant has been hypothesized to increase the amount of LRRK2 in an “off” state by reducing the amount of GTP-bound LRRK2 and increasing GTP hydrolysis, in contrast to effects observed with pathogenic variants in the ROC-COR domain of LRRK2 (*31, 37*). We show that the LRRK2 N551K R1398H variant reduces the levels of pT73 Rab10 on its own and in combination with disease-associated variants in the LRRK2 kinase domain using a combination of cellular models and PBMCs obtained from human carriers with and without protective haplotype. This data is in contrast to a previous report that showed no effect of this variant on pT73 Rab10 levels and may be explained by the more quantitative and sensitive assays and larger sample size employed in the current study (*31*). The LRRK2 protective haplotype was associated with reduced levels of total LRRK2 in cellular models, suggesting this variant might regulate the stability of LRRK2 in certain cell types. As the ROC-COR domain plays a key role in LRRK2 dimerization and the N551K R1398H variant did not impact LRRK2 levels in the presence of PD-associated variants in the ROC-COR domain, the protective variant may mediate its effects on LRRK2 levels by interfering with intermolecular interactions. Additional work is needed to better define how the protective LRRK2 haplotype leads to reduced kinase activity and to determine the consequences of these variants on the stability of LRRK2 itself. Together, our data suggest that the protective LRRK2 variants may be associated with lower PD risk by reducing LRRK2 activity and support the potential of therapeutic approaches that recapitulate these effects.

## Materials and Methods

### pT73 Rab10 monoclonal antibody generation

To generate antibody against phospho-T73-Rab10, peptide C-AGQERFH(pT)ITTSYYR (conjugated with either –KLH or -OVA) was used to inject 4 rabbits (New Zealand White rabbits) by Abcam. Rabbit bleeds/antisera were collected and screened by peptide ELISA assay and western blot. Two rabbits were selected for monoclonal antibody generation. Briefly, lymphocytes from rabbit spleen were fused with 240E-W2 cells to generate hybridomas. The multiclone hybridomas were screened by peptide ELISA, western blot and sandwich ELISA with cell lysate to identify antibodies for potential ELISA-based applications. 9 multiclones were selected and diluted to single clone. 18 monoclonal antibodies were further selected, and 8 of them were sequenced. The monoclonal antibody 19-4 was selected for pRab10 MSD assay.

### MSD assay

Capture antibodies were biotinylated using EZ-Link™ NHS-LC-LC-Biotin (Thermo Fisher, #21343), and detection antibodies were conjugated using Sulfo-TAG NHS-Ester (MSD, R31AA-1). 96-well (or 384-well) MSD GOLD Small Spot Streptavidin plates (MSD L45SSA-1) were coated with 25 μl (or 15 μl for 384 well plates) of capture antibody diluted in Diluent 100 (MSD, R50AA-2) for 1 hour at room temperature with 700 rpm shaking (1000 rpm for 384-well). After TBST wash (3X), 25 μl samples were added each well (10 μl for 384-well) and incubated at 4°C overnight with agitation at 700 rpm. After TBST wash (3X), 25 μl of detection antibodies (15 μl for 384-well) were added each well diluted in TBST containing 25% MSD blocker A (MSD R93AA-1), together with rabbit (Rockland Antibodies D610-1000) and mouse gamma globin fraction (D609-0100). After 1 hour incubation at room temperature at 700 rpm, followed by TBST washes (3X), 150 μl MSD read buffer (MSD R92TC, 1:1 diluted with water) is added (35 μl for 384-well), and plates are read on the MSD Sector S 600. For pT73 Rab10 MSD assays, all the three detection antibodies were tested with comparable performance. The detection antibody (Abcam, ab181367) was mainly used for data generation. GAPDH measurements were performed using the MSD assay kit (# K151PW).

### Antibodies used for the MSD assay

| Assay | Antibody type | targets | Vendor | Cat No. | Concentration<br>(μg/mL) |
| --- | --- | --- | --- | --- | --- |
| pS935<br>LRRK2 | Capture | pS935 LRRK2 | Abcam | ab133450 | 0.5 |
|  | Detection | Total LRRK2 | BioLegend | 808201 | 1 |
| Total LRRK2 | Capture | Total LRRK2 | BioLegend | 844401 | 0.5 |
|  | Detection | Total LRRK2 | BioLegend | 808201 | 1 |
| pT73 Rab10 | Capture | pT73 Rab10 | Denali | 19-4 | 1 |
|  | Detection | Total Rab10 | Abcam | ab181367 | 2 |
|  | Detection | Total Rab10 | Abcam | ab104859 | 2 |
|  | Detection | Total Rab10 | Creative<br>Diagnostics | DCABH-13141 | 2 |
| Total Rab10 | Capture | Total Rab10 | Creative<br>Diagnostics | DCABH-13141 | 1 |
|  | Detection | Total Rab10 | Abcam | ab181367 | 4 |

### MSD assay performance test with recombinant proteins, human PBMC and whole blood

The assay parameters (linear range and CV) were analyzed based on 384-well plates. Recombinant LRRK2 protein was from ThermoFisher (#A15197). For pT73 Rab10 assay, recombinant Rab10 protein (DU62271) and *in vitro* phosphorylated pT73-Rab10 protein (Estimated phosphorylation was 93% based on Phos-tag gel analysis) were from University of Dundee (https://mrcppureagents.dundee.ac.uk/). For total Rab10 assay, recombinant Rab10 protein (LSBio, LS-G2178) and Rab8 protein (LS-G29310) were used. Human cryopreserved PBMCs (AllCells, PB005F) were thawed and cultured in RPMI 1640 media, supplemented with 10% FBS and penicillin/streptomycin. Resuspended cells were filtered through 100 μm cell strainer, cultured in 96-well round bottom plates, and treated with MLi-2 (MedChemExpress LLC) for 1 hour. Frozen human whole blood samples were from Denali Employee Blood donation program and thawed and directly lysed in 96-well plates (100 μl whole blood sample with 100 μl lysis buffer). Samples were spun down (2,500x g for 20 minutes at 4°C) before MSD assay.

### Cell culture, transfection and treatment

*RAB8A* KO and *RAB10* KO A549 cells were generously provided by Dr. Dario Alessi (*11, 49*). HEK293 cells were transiently transfected with FLAG-tagged LRRK2 plasmids or HA-tagged Rab10 plasmid or mCherry-tagged Rab plasmids using Lipofectamine 3000, and lysed 48 hours after transfection. Cells were lysed in lysis buffer (Cell Signaling, #9803) supplemented with cOmplete tablet (Roche #04693159001), phosSTOP (Roche #04906837001) and Benzonase nuclease (Sigma, E1014).

### Western blot

The primary antibodies used were anti-pS935-LRRK2 (1:500, Abcam, ab133450), anti-LRRK2 (1:500, N241A/34, UC Davis/NIH NeuroMab Facility), anti-vinculin (1:500, Abcam, ab129002), anti-pT73 Rab10 (1:1000, Denali #19-4), anti-Rab10 (1:1000, Abcam, ab104859), anti-mCherry (1:1000, ab213511) and anti-actin (1:5000, Sigma, A2228). Details in Supplementary Methods.

### Animal model

All experiments in mice or rats were conducted in accordance with protocols approved by the Institutional Animal Care and Use Committee at Denali Therapeutics. Animals were housed under a 12-hour light/dark cycle and provided access to rodent chow and water ad libitum. LRRK2 G2019S KI mice (C57BL/6-Lrrk2tm4.1Arte) were maintained on a C57BL/6NTac genetic background at Taconic Biosciences. *LRRK2* KO rats (Long Evan, HsdSage:LE-*Lrrktm1Sage*) were obtained from Envigo.

### Animal tissue processing

Animals were deeply anesthetized with 2.5% Avertin (i.p.) and were transcardiac perfused with ice-cold PBS. Tissues were extracted, weighted and collected in 1.5 mL Eppendorf tubes, and immediately frozen on dry ice and stored at −80 °C. Tissues were homogenized in lysis buffer (10X of tissue weigh (mg) in μl) using the Qiagen TissueLyzer II for 2 × 3 minutes at 30Hz. Lung samples were lysed in CST (#9803) lysis buffer, and brain samples were lysed in RIPA buffer, and diluted 10 fold for MSD assay. Homogenates were spun at 14,000 rpm for 30 minutes at 4°C.

### iPSC-derived microglia cells

Human iPSCs (Thermo Fisher #A18945) were maintained in mTeSR1 medium (StemCell, #85857), and routinely passaged as clumps onto Geltrex (Thermo Fisher, #A1413301)-coated plates. Differentiation protocol was established previously (*25*). Briefly, iPSCs were first differentiated into hematopoietic progenitor cells (HPCs) using the STEMdiff Hematopoietic Kit (StemCell, #05310). HPCs that were positive for identity markers CD34, CD43, and CD45 were transferred to plates containing primary human astrocytes and co-cultured in Media C adapted from Pandya et al (*50*) for 14-21 days, during which time HPCs were progressively removed. The media included IMDM (Thermo Fisher), FBS, PenStrep, 20ng/mL each of IL3, GM-CSF, M-CSF (Peprotech). Once non-adherent cells in co-culture were predominantly mature microglia (>80%), the microglia were collected and plated for experiments.

### Generation of LRRK2 KO and N551K R1398H KI A549 cells with CRISPR

A549 cells were electroporated by AMAXA with guide-RNA Cas9-NLS ribonuclear protein (RNP) complexes pooled cells. The exon 1 of *LRRK2* gene was targeted to generate *LRRK2* KO cells, the guide 5’-tggctagtggcagctgtcagggg and 5’-agaaacgctggtccaaatcctgg were used. To introduce p. N551K (AAC > AAG) SNP modification, the guide ggtcctagcagctttgaaca was used. To introduce p. 1398H (CGT > CAT), the guide taatctttatttaggtcgtg was used. Individual clones were isolated from CRISPR pooled cells by limiting dilution. For LRRK2 KO cells, single clones were sequenced by PCR-TOPO cloning and further verified by western blot. For N551K R1398H KI cells, individual clones were screened and confirm by Sanger sequencing and next generation sequencing.

### LRRK2 and pRab10 measurement in healthy subjects and PD patients

Whole blood specimens from 12 PD patients (3 Female, 9 Male) and 6 Non-PD controls (2 Female, 4 Male) were obtained from the 24-hour Biofluid Sampling Study, in which biospecimens were donated by each subject at 11 time points over 26 hours (*51*). These specimens are a subset of a large Michael J. Fox Foundation biospecimen bank available to the community (https://www.michaeljfox.org/biospecimens).

### Isolation of PBMCs contributing to analysis of N551K R1398H relationship to LRRK2 and pRab10 levels

Subjects with and without a self-reported diagnosis of PD were recruited from the surrounding area as part of a biomarker study conducted from 2016 – 2017. The study was approved by a local Institutional Review Board and all subjects signed informed consents. Blood was collected into CPT-sodium heparin tubes (BD, BDAM362780) and PBMCs were then isolated following the manufacturer’s protocol.

### Statistical analysis of N551K R1398H relationship to LRRK2 and pRab10 levels

Samples used for the N551K R1398H genetic analysis were part of a dataset of 184 samples measured on the Illumina Infinium NeuroChip microarray platform(*52*). Standard processing, quality control, and imputation procedures were implemented (see the Supplementary Methods for details). Data for the LRRK2 SNPs G2019S (rs34637584), N551K(rs7308720), and R1398H (rs7133914) were extracted from the genotype data and used in subsequent association analysis. While the G2019S and N551K variants were ascertained directly by the NeuroChip array, the R1398H variant was imputed. We verified that this variant was imputed to a high quality (Minimac4(*53*), R-squared value > 0.99). Finally, we derived the N551K R1398H haplotype variable via the allele-level genotype data.

All statistical analysis was performed using R version 3.6.1. Of the 174 total samples with available imputed genotype data (see Supplementary Methods), 148 samples had complete data on the full set of covariates and MSD phenotypes used in association testing (see below) and were predicted to be of European ancestry. An additional 3 samples were removed due to outlying values (more than 3*SD from the mean across all samples for that phenotype) on 1 or more MSD phenotypes. Therefore, a total of 145 samples with joint genotype, MSD, and covariate data were available for association testing.

Prior to association testing, protein concentrations for total LRRK2, pRab10 and pS935 LRRK2 were normalized to GAPDH levels, and the pRab10/LRRK2 ratio was subsequently computed. These protein levels were then natural log-transformed and were fit in a linear model against sex, age, and the first two principal components (“Model 1”, see below) of the measured genotype array data (to account for genetic ancestry). The residuals were then inverse-normal transformed using the blom() function from the ‘rcompanion’ package (https://cran.r-project.org/web/packages/rcompanion/index.html). These transformed residuals were then used in association testing against the N551K R1398H haplotype. Because only 2 samples were homozygous haplotype carriers, we collapsed heterozygous and homozygous carriers together and tested for association in carriers (n = 15) relative to non-carriers (n = 130). Finally, we carried out sensitivity analyses to evaluate to the impact of disease status (“Model 2”), LRRK2 G2019S status (“Model 3”) or both (“Model 4”) on the statistical inferences from Model 1.

## Acknowledgments

We would like to thank Paul Davies and James Hastie at University of Dundee Medical Research Council Protein Phosphorylation Unit for providing phosphorylated Rab10 protein. We would also like to thank Dario Alessi at University of Dundee for generously sharing with us the *RAB8A* and *RAB10* KO A549 cells. The biospecimens from healthy subjects and PD patients from MJFF were kindly provided by the LRRK2 Biobanking Initiative site at the Columbia University Medical Center under the direction of Dr. Roy Alcalay. The LRRK2 Biobanking Initiative is coordinated and funded by The Michael J. Fox Foundation for Parkinson’s Research. We thank Janet Hurt, Dominique Jacquemet-Engelhart, Rinkal Chaudhary, and Anthony Lucas for their contributions to the PBMC isolation study contributing to the N551K R1398H findings. We also thank Joseph Lewcock, Anthony Estrada, Gilbert Di Paolo, Thomas Sandmann, and Steve Lianoglou for helpful feedback on the manuscript.

## Author contributions

XW, EN, MM, VB, SVA, MM, CL, HN, HS, AA, LP, NM, SHR and AGH designed experiments. XW, EN, RM, MM, VB, SVA, MM, CL, HN, HS and NM performed and analyzed experiments. XW, SVA, SHR and AGH wrote the manuscript.

## Data and materials availability

The data that support the findings of this study are available from the corresponding authors upon reasonable request.

