## Supplementary Information for "Understanding LRRK2 kinase activity in preclinical models and human subjects through quantitative analysis of LRRK2 and pRab10"

### Supplementary Materials

#### Supplementary Table 1

| Assay | LLOD (ng/mL) | LLOQ (ng/mL) | CV (%) | Linear range (ng/mL) |
|---|---|---|---|---|
| pS935 LRRK2 | 0.06 | 0.14 | 14.7 | 0.16-600 |
| LRRK2 | 0.03 | 0.07 | 15.8 | 0.06-96 |

Supplementary Table 1: pS935 LRRK2 and LRRK2 MSD assays with recombinant LRRK2 protein

#### Supplementary Table 2

| Assay | LLOD (ng/mL) | LLOQ (ng/mL) | CV (%) | Linear range (ng/mL) |
|---|---|---|---|---|
| pRab10 | 8.36 | 58.56 | 16.3 | 65.2-8350 |
| Rab10 | 0.02 | 0.07 | 11.8 | 0.56-45.7 |

Supplementary Table 2: pRab10 and Rab10 MSD assays with recombinant Rab10 protein

#### Supplementary Table 3

| Assay | LLOD (µg/mL) | LLOQ (µg/mL) | CV (%) | Linear range (µg/mL) |
|---|---|---|---|---|
| pS935 LRRK2 | 6 | 13.5 | 9.0 | 12~*3070 |
| LRRK2 | 3.2 | 6 | 7.5 | 12~*3070 |

*highest lysate concentration tested

Supplementary Table 3: LRRK2 MSD assay with human PBMC lysates

#### Supplementary Table 4

| Assay | LLOD (µg/mL) | LLOQ (µg/mL) | CV (%) | Linear range (µg/mL) |
|---|---|---|---|---|
| pRab10 | 48.5 | 191.9 | 12.4 | 191.9~*3070 |
| Rab10 | 8.5 | 24 | 9.8 | 3-12 |

*highest lysate concentration tested

Supplementary Table 4: pRab10 and Rab10 MSD assay with human PBMC lysates

### Supplementary Table 5

| Assay | LLOD (dilution factor) | LLOQ (dilution factor) | CV (%) | Linear range (dilution factor) |
|---|---|---|---|---|
| pS935 LRRK2 | 293 | 145 | 13.9 | 2-512 |
| LRRK2 | 512 | 295 | 9.2 | 2-512 |

Supplementary Table 5: LRRK2 MSD assay with human whole blood

### Supplementary Table 6

| | Model 1 | | | Model 2 | | | Model 3 | | | Model 4 | | |
|---|---|---|---|---|---|---|---|---|---|---|---|---|
| | Phenotype ~ Sex + Age + PC1 + PC2 | | | Phenotype ~ Sex + Age + PC1 + PC2 + Disease | | | Phenotype ~ Sex + Age + PC1 + PC2 + G2019S | | | Phenotype ~ Sex + Age + PC1 + PC2 + Disease + G2019S | | |
| Phenotype | Estimate | SE | P | Estimate | SE | P | Estimate | SE | P | Estimate | SE | P |
| total LRRK2 | 0.286 | 0.271 | 0.2918 | 0.217 | 0.271 | 0.4253 | 0.304 | 0.271 | 0.2628 | 0.21 | 0.271 | 0.4402 |
| pRab10 | -0.828 | 0.263 | 0.002 | -0.806 | 0.263 | 0.0026 | -0.834 | 0.263 | 0.0018 | -0.792 | 0.264 | 0.0031 |
| pS935 LRRK2 | 0.127 | 0.271 | 0.6407 | 0.025 | 0.272 | 0.9272 | 0.117 | 0.272 | 0.6675 | 0.003 | 0.272 | 0.9903 |
| pRab10/LRRK2 | -0.697 | 0.265 | 0.0096 | -0.62 | 0.267 | 0.0216 | -0.676 | 0.266 | 0.012 | -0.587 | 0.267 | 0.0296 |

Supplementary Table 6: Linear regression statistics for association testing of LRRK2 and pRab10 levels against N551K R1398H haplotype status.

Protein levels were normalized to GAPDH levels, natural log transformed, and then fit in a linear model against the covariates listed under each "Model". Residuals from each model were the inverse normal transformed and tested for association against the N551K R1398H haplotype variable. Estimate = standard deviation change in adjusted and transformed protein level for haplotype carriers relative to non-carriers. SE = standard error of estimate.

**Supplementary Figure S1**

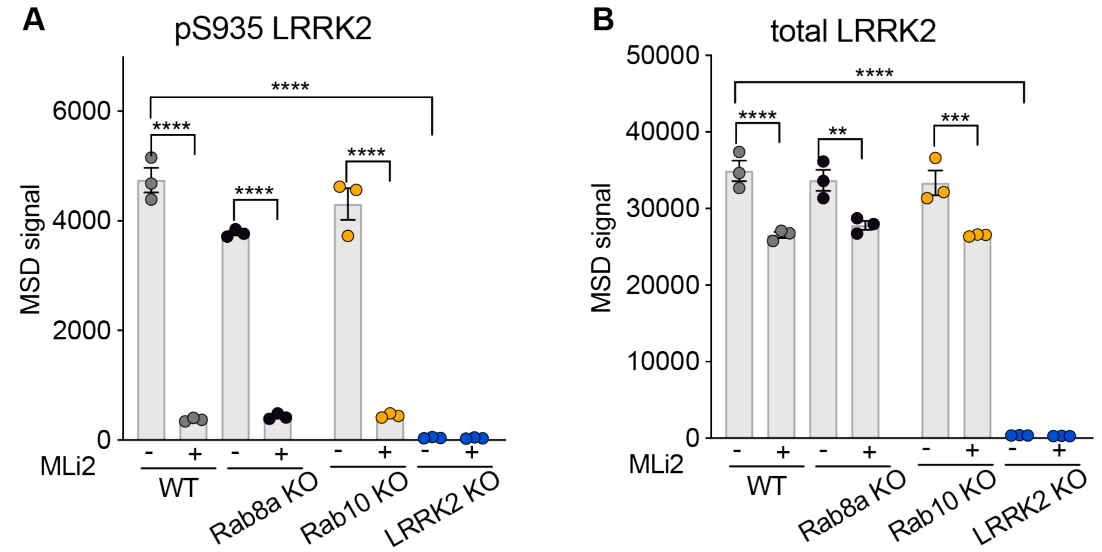


Supplementary Figure S1. pS935 LRRK2 and total LRRK2 levels measured by MSD in A549 cells of different genetic backgrounds.

**A-B)** pS935 LRRK2 and total LRRK2 levels were comparable in *RAB8A* or *RAB10* KO A549 cells and were absent in *LRRK2* KO cells. Data shown as mean ± SEM with p values: one-way ANOVA with Sidak's multiple comparison test. ** $p \leq 0.01$, *** $p \leq 0.001$, **** $p \leq 0.0001$.

**Supplementary Figure S2**

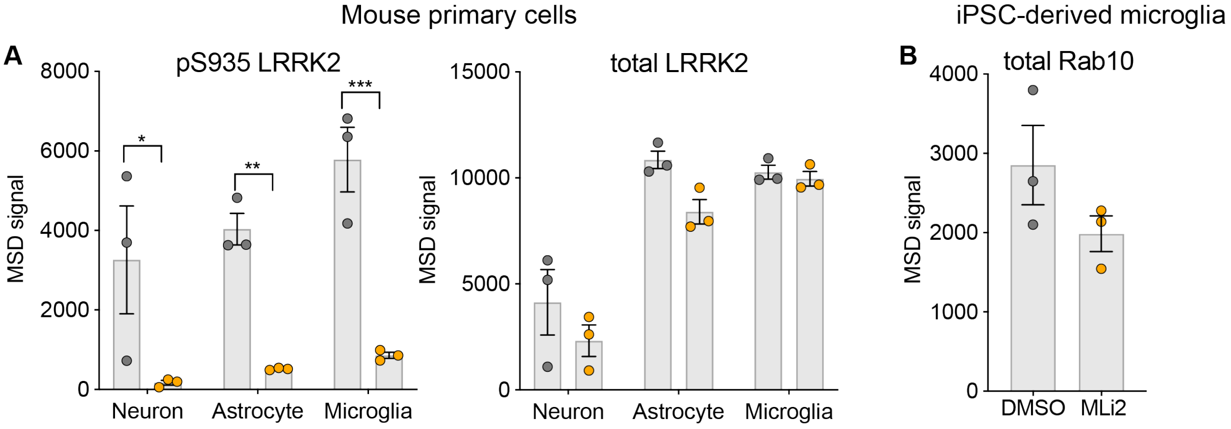


Supplementary Figure S2. LRRK2 is highly expressed in mouse glia cells

**A)** LRRK2 is highly expressed in mouse primary cultured cortical astrocytes and microglia, and inhibition of LRRK2 kinase by MLi-2 (500 nM, 2 hours) significantly reduced pS935 LRRK2. pS935 and LRRK2 levels were assessed by MSD assay. n=3. Data shown as mean ± SEM with p values: two-way ANOVA with Sidak's multiple comparison test. **B)** No significant difference in Rab10 levels with and without MLi-2 in iPSC-derived microglia. n=3, Data shown as mean ± SEM. Paired t-test. * $p \leq 0.05$, *** $p \leq 0.001$

**Supplementary Figure S3**

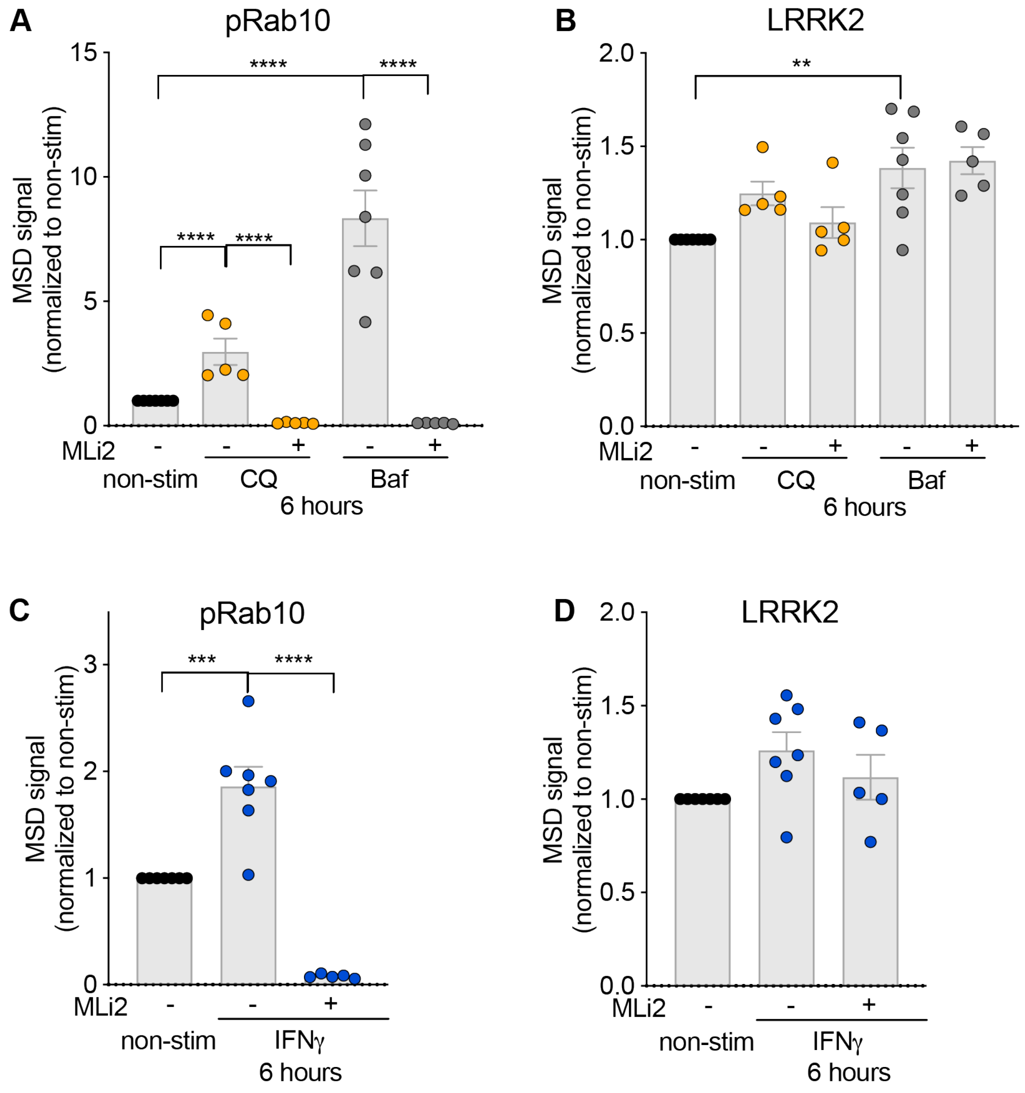


Supplementary Figure S3. Lysosomal stressors and inflammatory stimuli modulate LRRK2 levels and activity in iMG.

**A-B)** In iMicroglia, acute treatment of chloroquine (50 mM, 6 hours) and bafilomycin A1 (100 nM, 6 hours) increased pRab10 level, which can be blocked by LRRK2 kinase inhibitor (MLi-2, 500 nM) treatment. LRRK2 levels were largely not affected by the lysosomal stressors, and only with a mild increase after Bafilomycin A1 treatment. **C-D)** Acute IFN- $\gamma$ treatment (20 ng/mL, 6 hours) induced a significant increase in pRab10 levels in iMicroglia cells, with minimal effects on LRRK2 levels. n=5-8. Data shown as mean ± SEM with p values: one-way ANOVA with Sidak's multiple comparison test. ** $p \leq 0.01$, *** $p \leq 0.001$, **** $p \leq 0.0001$.

### Supplementary Figure S4

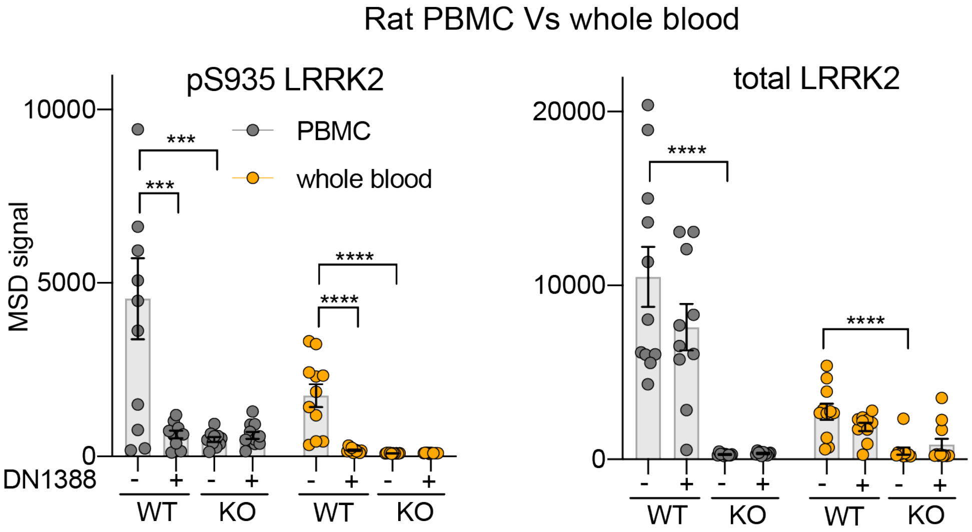


Supplementary Figure S4

pS935 LRRK2 and LRRK2 can be specifically measured in rat PBMC and whole blood from WT and *LRRK2* KO rats with and without LRRK2 kinase inhibitor (DN1388, 100 mg/kg, QD, PO dosing for 10 days). n=11 animals/group. Data shown as mean ± SEM with p values: two-way ANOVA with Tukey's multiple comparison test. *** p≤ 0.001, **** p≤ 0.0001.

### Supplementary Figure S5

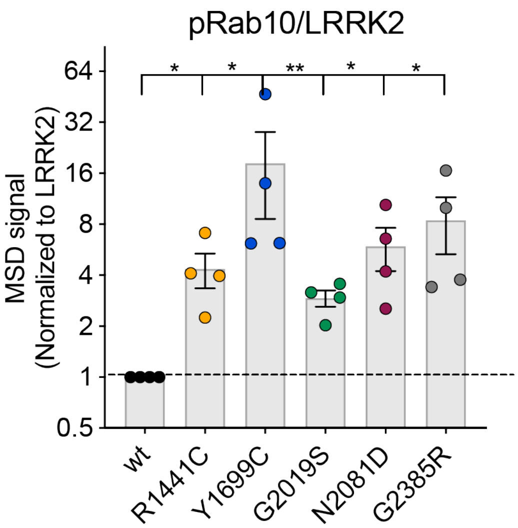


Supplementary Figure S5.

Expressions of PD and CD risk variants increase pRab10 level (normalized to LRRK2) in HEK293T cells overexpressing LRRK2 and Rab10. n=4. Data shown as mean ± SEM with p values: one-way ANOVA with Dunnett's multiple comparison test. * $p \leq 0.05$, ** $p \leq 0.01$.

### Supplementary methods

*Western blot*

Cell or tissue lysates were normalized for equal protein loading and were prepared by incubating with NuPage LDS Sample Buffer (ThermoFisher, NP0007) and NuPAGE™ Sample Reducing Agent (ThermoFisher, NP0004) for 10 min at 70 °C to denature samples. Lysates were loaded onto NuPAGE 4-12% Bis-Tris gels (Invitrogen). Proteins were transferred to nitrocellulose membranes for 7 min (Trans-Blot Turbo Transfer System, Bio-Rad). Membranes were blocked with Rockland blocking buffer, incubated with primary antibody overnight at 4 °C, and then with secondary antibodies (1:20,000, LI-COR) for 1 hour at room temperature. LI-COR Odyssey system was used for Western blot detection and quantitation.

*Genotype array data measurement, processing, and quality control*

Samples used for the N551K R1398H genetic analysis were part of a dataset of 184 samples measured on the Illumina Infinium NeuroChip microarray platform (*52*). Whole genome amplification of DNA extracted from PBMCs, DNA fragmentation, two-step allele detection involving hybridization and single base extension were performed according to the manufacturer's instructions. Briefly, whole genome amplification occurred at 37°C for 20-24 hours, followed by enzymatic fragmentation at 37°C for 1 hour. The DNA was purified by alcohol precipitation, resuspended and hybridized on Illumina BeadChips at 48 °C for 16-24 hours. After single base extension and fluorescent labeling, the BeadChips were scanned using an Illumina iScan. Genotypes were called using the Neuro_Consortium_v1-1_20015375_A1.bpm annotation file (GRCh37 / hg19) via Illumina's GenomeStudio software. All steps from DNA extraction through genotype calling were performed by Q2 Solutions (Morrisville, NC, United States).

Prior to SNP-level and sample-level quality control carried out in Plink v1.9(*54*) https://www.cog-genomics.org/plink/1.9/), genotype array data consisted of 487,374 variants measured on 184 samples. SNPs were removed if their Hardy-Weinberg Equilibrium p-value (estimated using control samples only) was < 1E-5, if their minor-allele frequency (MAF) was less than 0.01, or if their call failure rate exceeded 5% of samples. Samples were removed if their call failure rate exceeded 5% of variants, and if they had outlying rates of heterozygosity (more or less than 3 SD from mean rate across all samples). To account for relatedness, we retained the sample with lower rate of genotype missingness among any pairs of samples in which the proportion of identity-by-descent metric (determined from LD-pruned data) exceeded 0.2. After these QC steps, genotype data was available on 302,591 variants and 174 samples.

Next we used the imputation preparation tool available from the McCarthy Group (https://www.well.ox.ac.uk/~wrayner/tools/) to perform strand flipping, reference/alternate allele assignment and other variant-level data checks prior to merging with 1000 Genomes (1000G) Phase 3 reference samples (*55*). We performed principal component analysis (PCA) on this merged dataset and used the first 10 principal components and 1000G super population labels to predict genetic ancestry for all samples in our dataset via the k-nearest neighbors algorithm. We then performed phasing using Eagle v2.4.1(*56*) and imputed the data using reference panels from the predicted ancestry of the study samples. Imputation was performed using Minimac4(*53*) with default parameters, and DosageConvertor (https://genome.sph.umich.edu/wiki/DosageConvertor) was used to convert Minimac4 output to Plink dosage format. Finally Plink v2(*54*) https://www.cog-genomics.org/plink/2.0/) was used to convert the data to hard calls under the default parameters.
